# A Cross-Sectional Study of Prevalence and Spatial Patterns of Major Limb Loss in the Acholi Sub-Region of Uganda

**DOI:** 10.1101/2020.05.14.095836

**Authors:** Pamela Atim, Constantine S. Loum, Tom R. Okello, Samuel M. Magada, Walter Onen Yagos, Peter Abelle, Emmanuel B. Moro, Jonathan J. Huck, Anthony Redmond, Mahesh Nirmalan

## Abstract

**BACKGROUND:** There is a widely reported preponderance of major limb loss in Northern Uganda, which is believed to have been caused in large part by prolonged civil war. Access to rehabilitation facilities is extremely limited, and there has never been a study to understand how many people have major limb loss, nor how many of them have had access to medical or rehabilitative services.

**AIM:** The first prevalence study of disability and major limb loss in the region, and evaluated spatial patterns of cases of MLL.

**DESIGN:** Cross-sectional survey.

**SETTING:** This research was undertaken in a community setting (at subjects’ homes).

**POPULATION:** 7,864 randomly selected households throughout the Acholi Sub-Region of Northern Uganda.

**METHODS:** This study comprised two questionnaires, the first to be completed by the head of every sampled household (*n*=7,864), and the second by any member of the household with major limb loss (*n*=181). Household locations were examined for spatial autocorrelation using Moran’s *I* statistic. The χ^2^ goodness of fit statistic was used to profile those with major limb loss in comparison with the underlying population.

**RESULTS:** We conservatively estimate that there are c.10,117 people with major limb loss in the region who require long-term rehabilitation services (c.0.5% of the population), and c.150,512 people with disabilities other than MLL (c.8.2% of the population). People with major limb loss are spread throughout the region (as opposed to clustered in specific locations) and are disproportionately male, older and less well educated than the general population.

**CONCLUSION:** This research demonstrates, for the first time, the extent of the inadequacy of long-term rehabilitation services for those with major limb-loss in the study area. We provide new insight into the reasons that people are not accessing medical and rehabilitative services, and propose a way forward through the successful demonstration of an ‘outreach clinic’ model.

**CLINICAL REHABILITATION IMPACT:** The discovery of the spatial pattern of those with major limb loss, alongside the demonstration of the clinical outreach model, provides a compelling argument for the need of more such services and associated policy frameworks in remote and rural regions in the Global South.

**TIDIER Checklist:** We confirm that this manuscript conforms to the STROBE Checklist for Cross-Sectional Studies

## Introduction

### Background

The Acholi Sub Region in Northern Uganda is a relatively impoverished part of the world which suffered a prolonged war which lasted for over 20 years. In this conflict, a rebel organisation known as the Lord’s Resistance Army (LRA) fought the Ugandan Government forces, resulting in large scale destruction of property, mass displacement and extensive physical injuries to the civilian and combatant populations due to explosions, gun short and punishment amputations [1–3]. The extensive media coverage associated with this conflict triggered large-scale operations by relief agencies such as the United Nations, World Food Programme, International Committee of the Red Cross, AVSI, and several other smaller International Non-Governmental Organisations (INGOs). These voluntary agencies accounted for the vast majority of health provision during the conflict, as the already limited local health infrastructure was destroyed by war [4]. However, these services were not sufficient to meet demand, and were often not able to reach the most remote and rural communities, leaving many un-treated. To compound these issues, the involvement of these agencies has gradually scaled down since the end of the conflict in 2005. Limited resources and the remote nature of this region means that adequate capacity for rehabilitative service provision has not been created in the intervening period and many individuals have never received treatment and necessary assistive devices thus maximising any residual potential.

The significant health benefits associated with the provision of a suitable prosthesis for people with major limb loss (MLL) are widely understood [5, 6], but difficulties in accessing such services are well recognised in Uganda and many other countries in Sub Saharan Africa. The relatively low priority given to resource provision for less visible chronic health needs such as rehabilitation services is very apparent in many countries, and Uganda (Northern Uganda in particular) is no exception to this pattern [7]. Despite the formation of positive government policy with respect to disabled people, the disability services of the Ugandan government suffers from differing statistics and of definitions of disability, both of which make targeting interventions difficult [8, 9]. Data on disability type and prevalence are also scarce, mostly existing only at the national level, with no understanding of spatial variations of prevalence within Uganda [8]. Such detailed local data are of particular importance in a country where a prolonged and violent conflict was concentrated in some of the most remote parts of the country, where data collection is likely to be worst. In this context, prevalence studies are essential tools to highlight the magnitude of the problem and its potential impact on wellbeing and productivity in order to influence health policy and establish the scale of rehabilitation service requirements in regions such as this.

### Objectives

The present study therefore seeks to address this deficit by systematically determining the prevalence and spatial distribution of Major Limb Loss (MLL) in the Acholi sub-region of Northern Uganda. The study seeks to:

1. Characterise the types and causes of major disability and MLL in the population in Northern Uganda
2. Provide an estimate for the number of people with MLL in the Acholi Sub-Region
3. Describe the demographic characteristics of those with MLL in the Acholi Sub-Region
4. Characterise the spatial distribution of people with MLL in the Acholi Sub-Region

## Materials and Methods

### Study Area

This work is focused upon the Acholi sub-region of Northern Uganda, which comprises eight districts: Agago, Amuru, Gulu, Kitgum, Lamwo, Nwoya, Omoro and Pader. The region is approximately 28,500 km^2^ in area [10], and the total population of this area is currently estimated at approximately 1,790,700 (excluding refugees) [11], plus approximately 55,574 refugees [12]; approximately 1,846,274 in total. However, the population of this region is extremely difficult to determine accurately, in part due to the remote and rural nature of the region, significant unquantified uncertainty in the Ugandan census (which is based on sampling rather than comprehensive survey), extremely rapid population growth, and frequent movement of people (including refugees) in both directions across international borders to the Democratic Republic of the Congo and South-Sudan.

### Study Design

Our study is based on a cross-sectional survey of randomly selected clusters of private household units selected from across the entire region. Our initial power calculations indicated that it would be necessary to study a random sample of 16,497 people in order to achieve a confidence level of 99% with a margin of error of 1%, which equates to approximately 3,000 households [13], given an average household size for Acholi of 5.5 people [14]. However, in this study adopted a deliberate strategy of oversampling in order to account for the high degree of uncertainty in the population estimate; the expectation of low prevalence of MLL; and the expectation of spatial autocorrelation (clustering) in MLL cases. We therefore set out to survey 8,000 households across the 8 districts, which equates to c.44,000 people [14].

The sample of 8,000 households was selected using a two-stage random sampling approach using GIS (Geographical Information Science). In the first stage, a stratified approach was used in which 100 locations were selected within each of the eight districts. In the absence of up-to-date maps or data on the distribution of the population in the region [15], we utilised the High-Resolution Settlement Layer (HRSL) dataset [16], which was produced in 2015 through collaboration between the World Bank and the Connectivity Lab at Facebook [17, 18]. This dataset provides estimated values of population density as a continuous raster surface of *c*.30m^2^ (1 arc-second^1^) cells. Values in the HRSL are calculated using high-resolution satellite imagery and a combination of computer vision and machine learning techniques in order to detect buildings and combine this information with census data in order to determine a population estimate [17, 18]. In order to select the sample locations, all cells containing non-zero values in the HRSL dataset were converted to point locations using a Python script [19], and then 100 per district (800 in total) were selected using the ‘Random Selection’ functionality in QGIS [20]. The distribution of the 800 sampling locations is shown in Figure 1, along with an illustration of the straight-line distance to the Gulu Referral Orthopaedic Clinic, which is the only facility in the region for MLL rehabilitation and the provision of prosthetic limbs. In the second stage, at each of the 800 randomly selected sample locations, the closest 10 households were identified for further assessment, resulting in a sample of 8,000 households.

**Figure 1:**
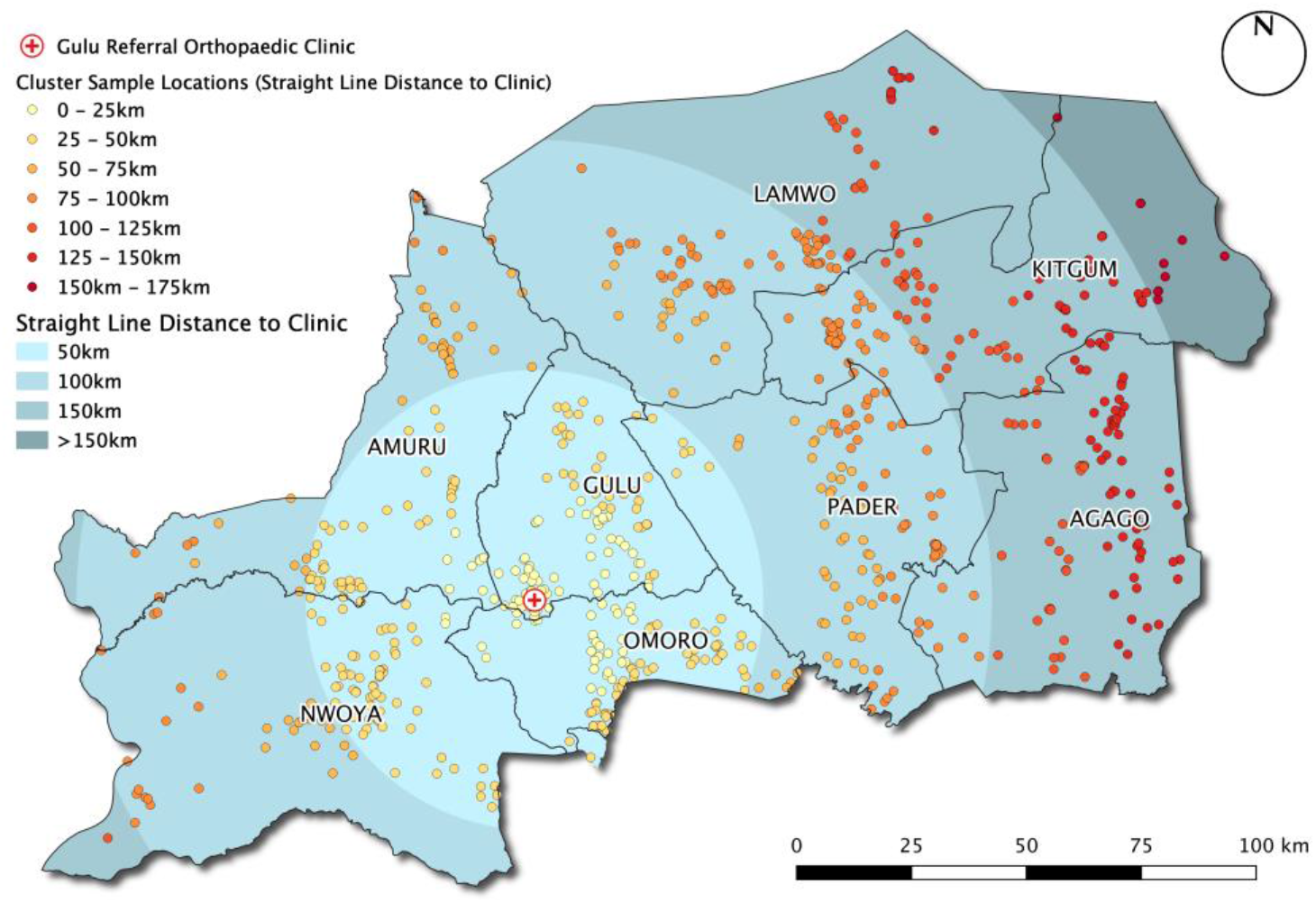
The distribution of the 800 sample locations across the entire region consisting of 8 districts

At each household, the study comprised two pre-tested, researcher-administered semi-structured questionnaires. The first questionnaire (S1: Appendix 1) was completed as part of an interview of the Head of Each Household (HoH) at each of the 8,000 households. The second questionnaire (S2: Appendix 2) was completed as part of an interview of each person identified to have MLL within those households. In both cases, the only exclusion criteria were that the consenting participant had to be able to understand the purpose of the research in order to give informed consent, and be over 18.

Both questionnaires (S1 and S2) were developed through multi-stakeholder engagement including disabled people attending the regional rehabilitation centre at the Gulu Referral Hospital, community health workers and academics from the Gulu Medical Faculty. The validity of the questionnaire was tested amongst health workers representing the Gulu Regional health office, volunteers representing the Gulu Medical Faculty and a cohort of disabled people attending the Gulu Referral Hospital. Through these pre-evaluations and focus group discussions the questions were refined in an iterative fashion in order to ensure reproducibility and clarity thereby ensuring the ability of the field teams to administer the survey tools effectively. Both sets of survey tools were then translated to the local Acholi Language by trained translators and the validation process was repeated for the translated survey tools with iterative corrections when needed. The validation studies for the survey tools were conducted in May 2018.

### Data Collection

The field surveys were conducted between July and September 2018, during which time a team of researchers visited all 8,000 selected sample locations. Trained research assistants administered both questionnaires during a 1:1 interview with each participant, which was necessary due to low levels of literacy in the region. All participants provided informed written consent prior to being interviewed. Interviews were conducted in the local language (Acholi) and translated into English for analysis. The local village administration officer (known as the ‘Local Councillor One’, or ‘LC1’) was given prior notice of the research team’s visit to any given village and was present alongside the visiting research teams in accordance with cultural norms.

The frequency and causes for MLL within the sample population were calculated, and the observed gender, age and educational attainment distributions were compared against the expected values for this population obtained from national census data using the Chi-squared (χ^2^) goodness of fit statistic. Expected distributions for this comparison were determined by scaling the distribution of values drawn from the UBOS 2014 Census [21] or UBOS 2018 Mid-Year Population Projection [22]. In order to examine spatial distribution of MLL across the Acholi Sub-Region, we used the *Moran’s I* statistic [23] in order to test for spatial autocorrelation (in which the null hypothesis is that similar data points are randomly dispersed). Such analysis allows us to establish the extent to which areas with high or low incidence of MLL are located closer to each other (spatial clustering) or further apart from each other (spatial dispersal), than would be expected. This statistic is calculated for MLL cases aggregated into sub-counties, using queen’s weighting (in which polygons are considered adjacent if they share at least one vertex). Significance of the finding is calculated as a pseudo-*p* value, based on 9999 permutations. No attempt has been made to adjust values for relative population of each sub-county because this is already accounted for in the design of the sub-counties. Statistical analysis was undertaken using Python 3.8.

This research was approved by the following ethics boards: The University of Manchester Research Ethics Committee (2017-2624-3985); Gulu University Research Ethics Committee (GUREC-033-18); Uganda National Council for Science and Technology (SS4817).

## Results

### The Types and Causes of Disability and MLL in Acholi

A total of 8,000 households were visited by researchers, of which 7,864 HoH responses were available for analysis following data cleaning. 3,763 people with disabilities were identified in the sample households (47.9%), comprising 3,943 individual disabilities (180 individuals had multiple disabilities). Types and causes of disability data are summarised in Table I.

**Table I:**
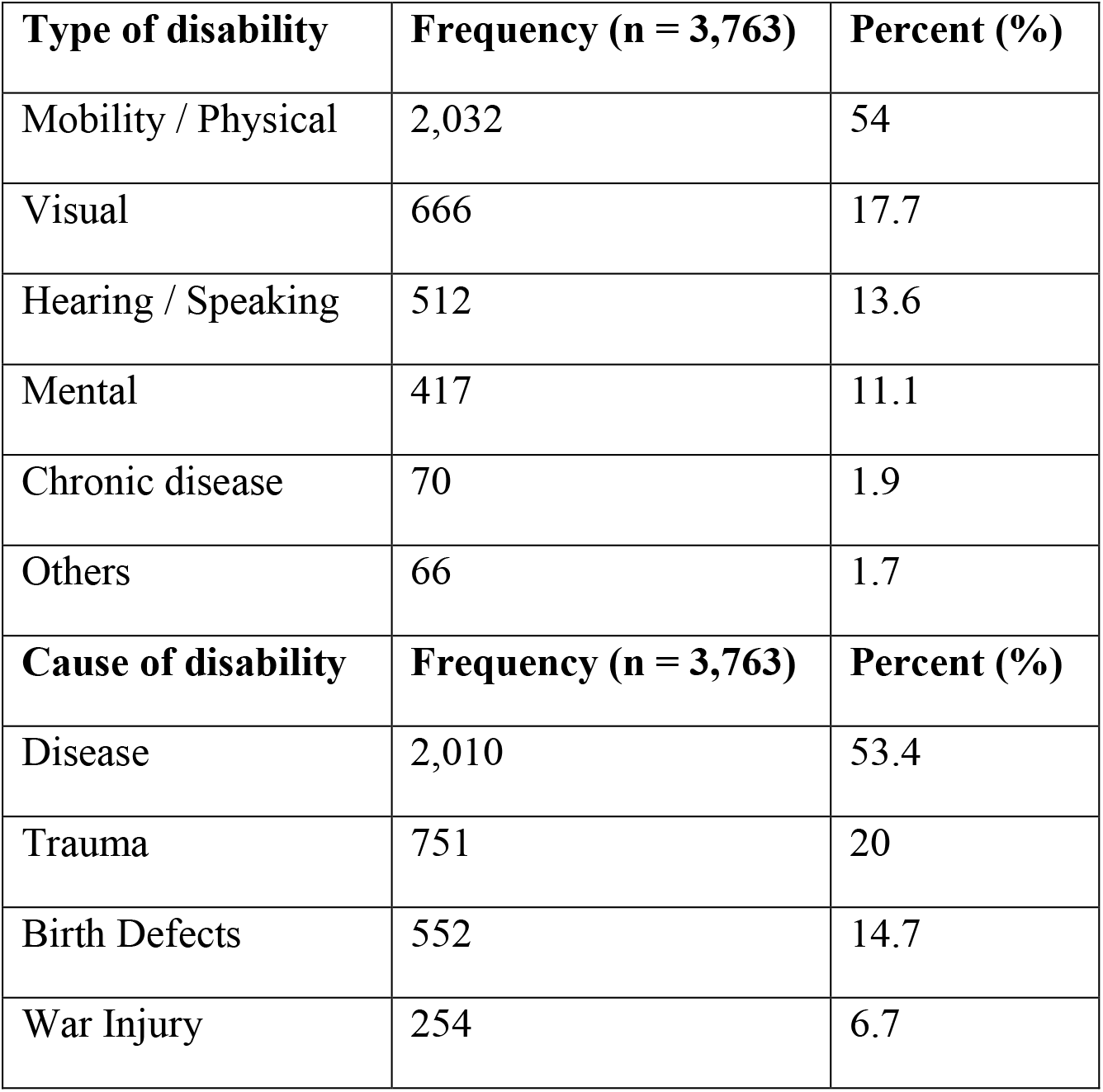
Disabilities by cause and type

| **Type of disability** | **Frequency (n = 3,763)** | **Percent (%)** |
| --- | --- | --- |
| Mobility / Physical | 2,032 | 54 |
| Visual | 666 | 17.7 |
| Hearing / Speaking | 512 | 13.6 |
| Mental | 417 | 11.1 |
| Chronic disease | 70 | 1.9 |
| Others | 66 | 1.7 |
| **Cause of disability** | **Frequency (n = 3,763)** | **Percent (%)** |
| Disease | 2,010 | 53.4 |
| Trauma | 751 | 20 |
| Birth Defects | 552 | 14.7 |
| War Injury | 254 | 6.7 |

Of the 3,763 individuals with disabilities, 237 had MLL (c.6%); of which c.59% had lost lower limbs, c.39% upper limbs, and c.2% both upper and lower limbs. Approximately 98% of people with MLL had unilateral limb loss, whilst c.2% had bilateral limb loss. These data are summarised in Table II. We were able to administer the second questionnaire to 181 of the 237 people with MLL (c.76%). Reasons for non-participation in both questionnaires were limited to the exclusion criteria described above.

**Table II:**
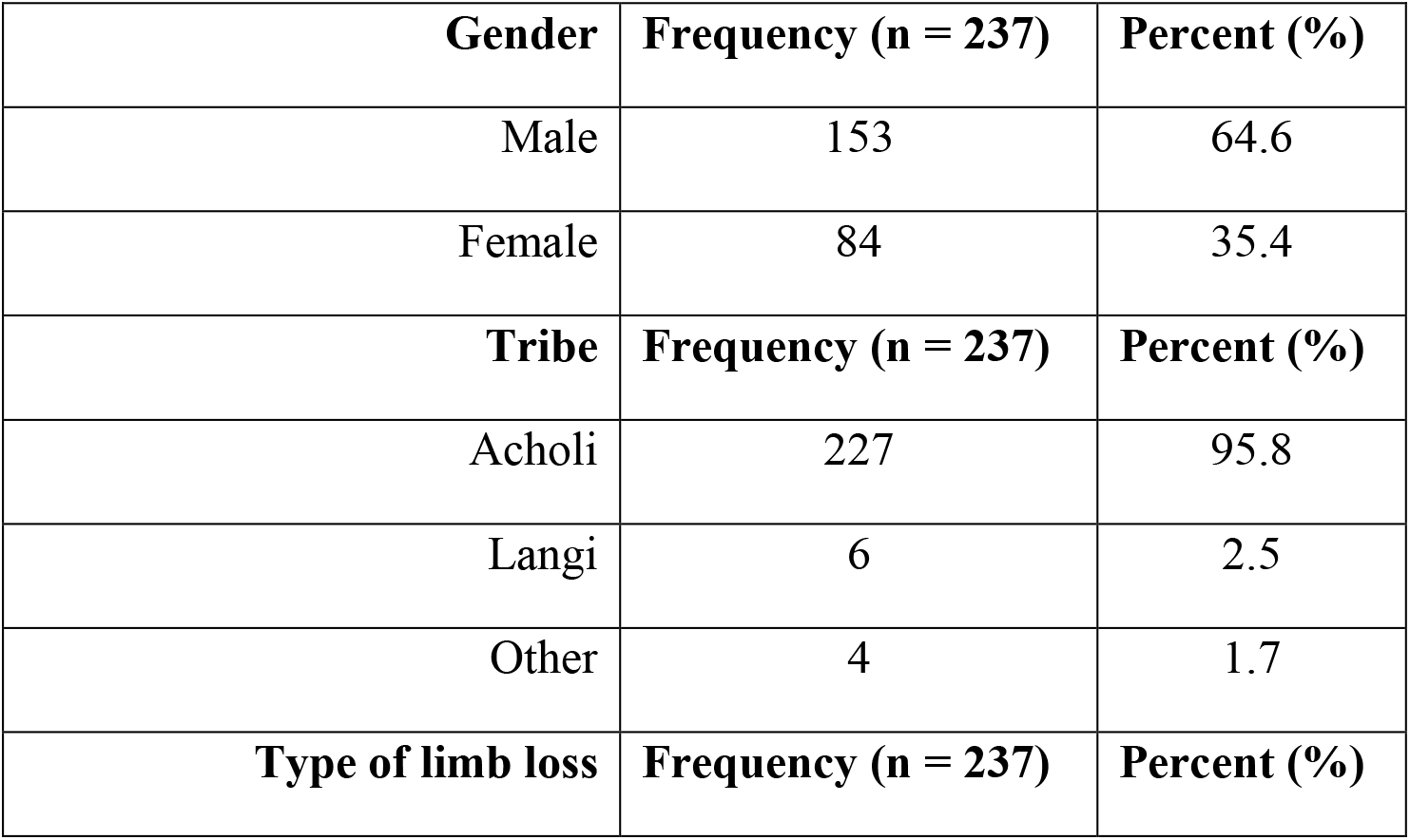
Characterising major limb loss

| **Gender** | **Frequency (n = 237)** | **Percent (%)** |
| --- | --- | --- |
| Male | 153 | 64.6 |
| Female | 84 | 35.4 |
| **Tribe** | **Frequency (n = 237)** | **Percent (%)** |
| Acholi | 227 | 95.8 |
| Langi | 6 | 2.5 |
| Other | 4 | 1.7 |
| **Type of limb loss** | **Frequency (n = 237)** | **Percent (%)** |
| Lower limbs | 139 | 58.7 |
| Upper limbs | 93 | 39.2 |
| Upper and lower limbs | 5 | 2.1 |
| **Level of limb loss** | **Frequency (n = 237)** | **Percent (%)** |
| Above knee joint (proximal) | 87 | 36.8 |
| Below knee joint (distal) | 47 | 19.8 |
| Above elbow joint (proximal) | 43 | 18.1 |
| Below elbow joint (distal) | 60 | 25.3 |

The main causes of limb loss are illustrated in Figure 2. As expected, war injuries are the most frequent cause of MLL in the region, accounting for c.48.9% of cases. Amongst the MLL cohort 46% had not had access to assistive devices (prosthetic limbs, crutches, walking stick etc.); 47% had no access to any rehabilitative services; and 10% had no access to any kind of healthcare whatsoever (including through local healthcare volunteers).

**Figure 2:**
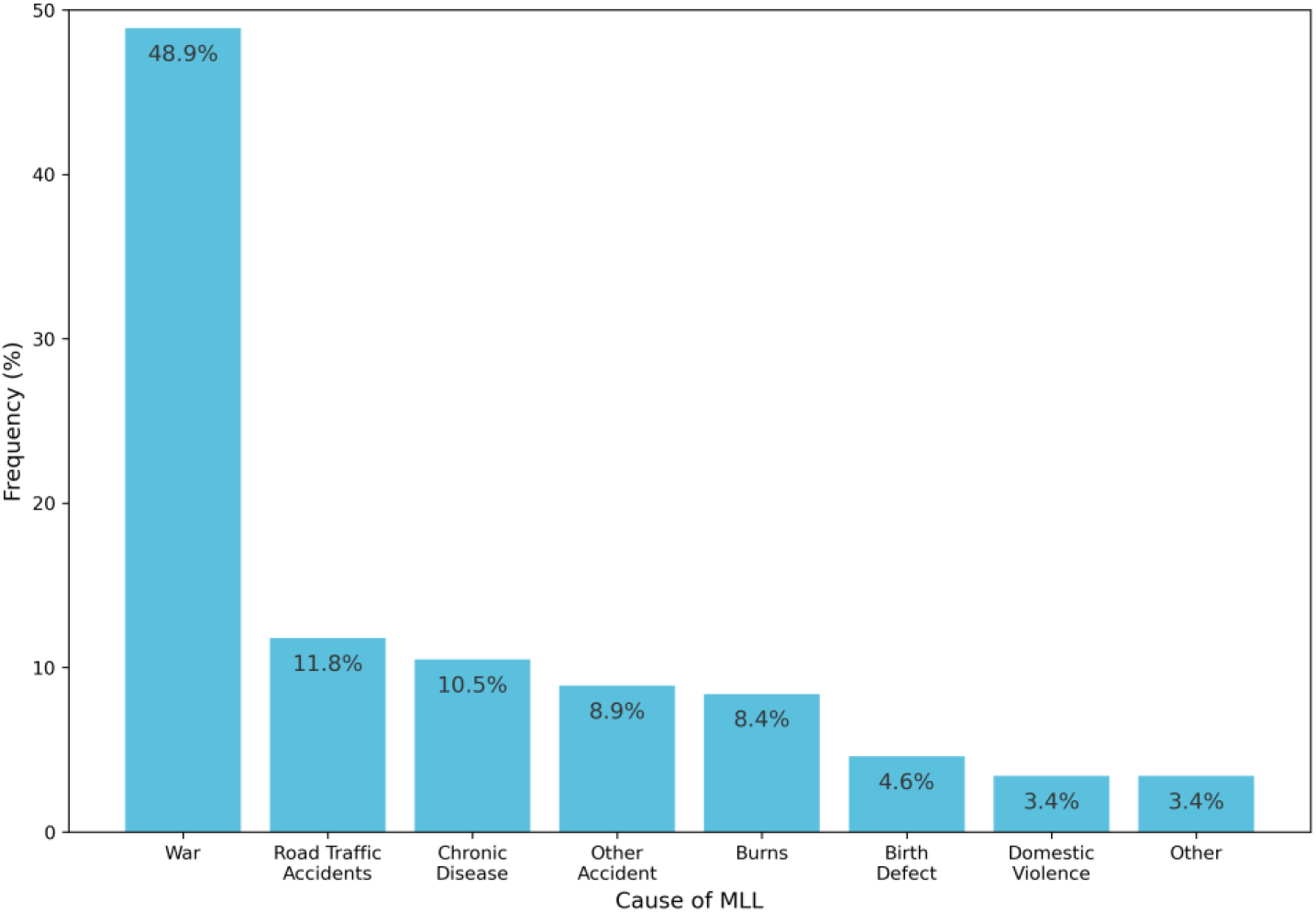
Aetiology of Major Limb Loss (MLL)

### Estimating Disability and MLL Prevalence in Acholi

The mean household size in the Acholi Sub-Region is 5.5 [14], so our 7,864 households equate to approximately 43,252 people, of which 237 had MLL. Because the sample was randomised, we can scale this up to the population of the Acholi Sub-Region using a conservative population estimate of 1,846,274 people [11, 12]. Based upon this, we estimate that there are approximately 10,117 people with MLL in the Acholi Sub Region (c.0.5% of the population), which is the first formal estimate of MLL prevalence in the region. Similarly, we estimate that there are approximately 150,512 people with disabilities other than MLL in the region (c.8.2% of the population), many of whom once again have had little or no access to medical or rehabilitation services. Given the aforementioned expectation that the population projections used in this study are likely to be a substantial underestimate, these figures should in turn be considered as conservative estimates.

### Socio-Demographic characterisation of those with MLL in Acholi

The gender, age, and education of those with MLL was compared with underlying population distributions using the χ^2^ goodness of fit statistic, in order to identify any socio-demographic trends amongst those who are disabled. Of the 181 participants that completed the questionnaire, gender and age data were available for 167, and education was available for 166; missing data were disregarded for the purposes of this analysis (Figure 3a–c). This analysis suggested that those with MLL in the Acholi Sub-Region are disproportionately male (*p*=3.1×10^−6^), older (*p*=1.3×10^−67^) and less well educated (*p*=8.9×10^−73^) than would be expected from the respective underlying population distributions. The expected and observed distributions (as well as their differences) as illustrated in Figure 3a, 3b and 3c for gender, age and education respectively. In all three cases, the probability that the distribution of observed values could have been drawn from the expected population distribution is vanishingly small (all are significant at the 99.9% confidence level).

**Figures 3a, 3B,3C & 3D:**
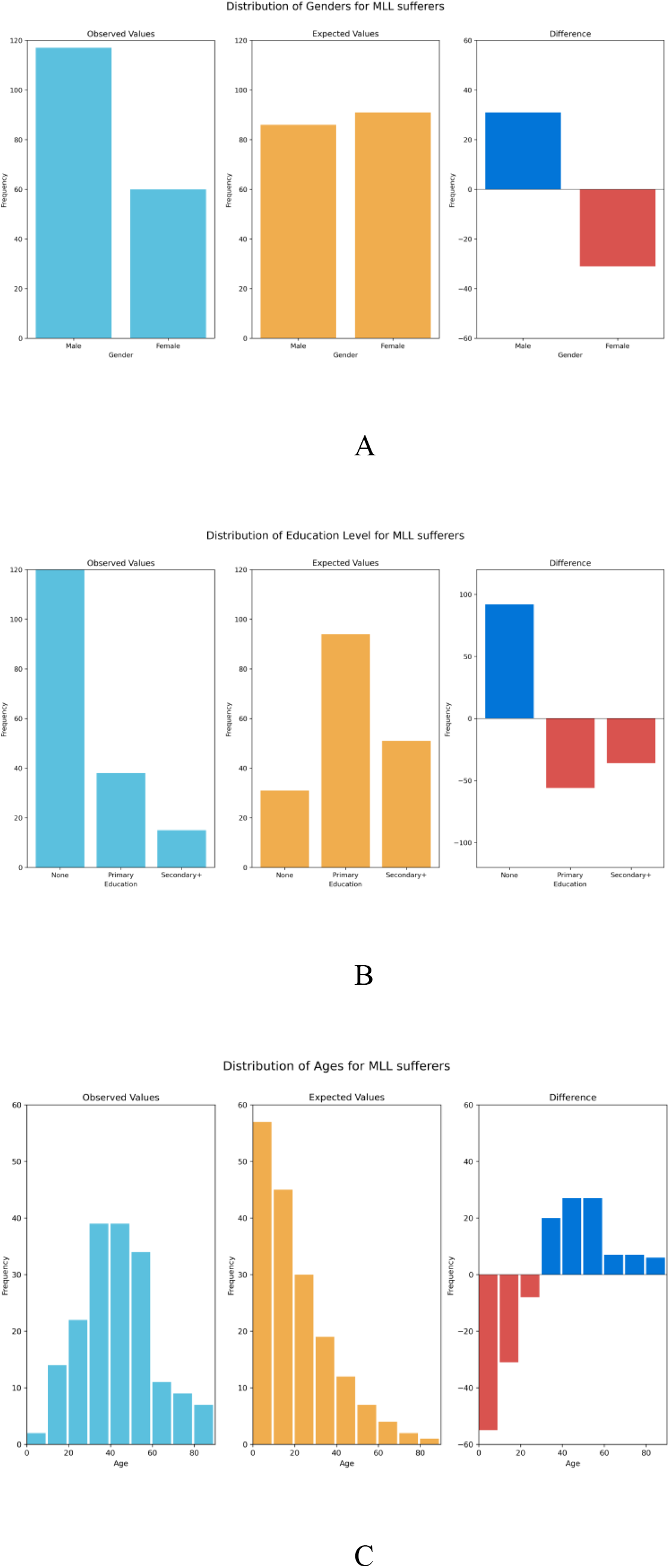
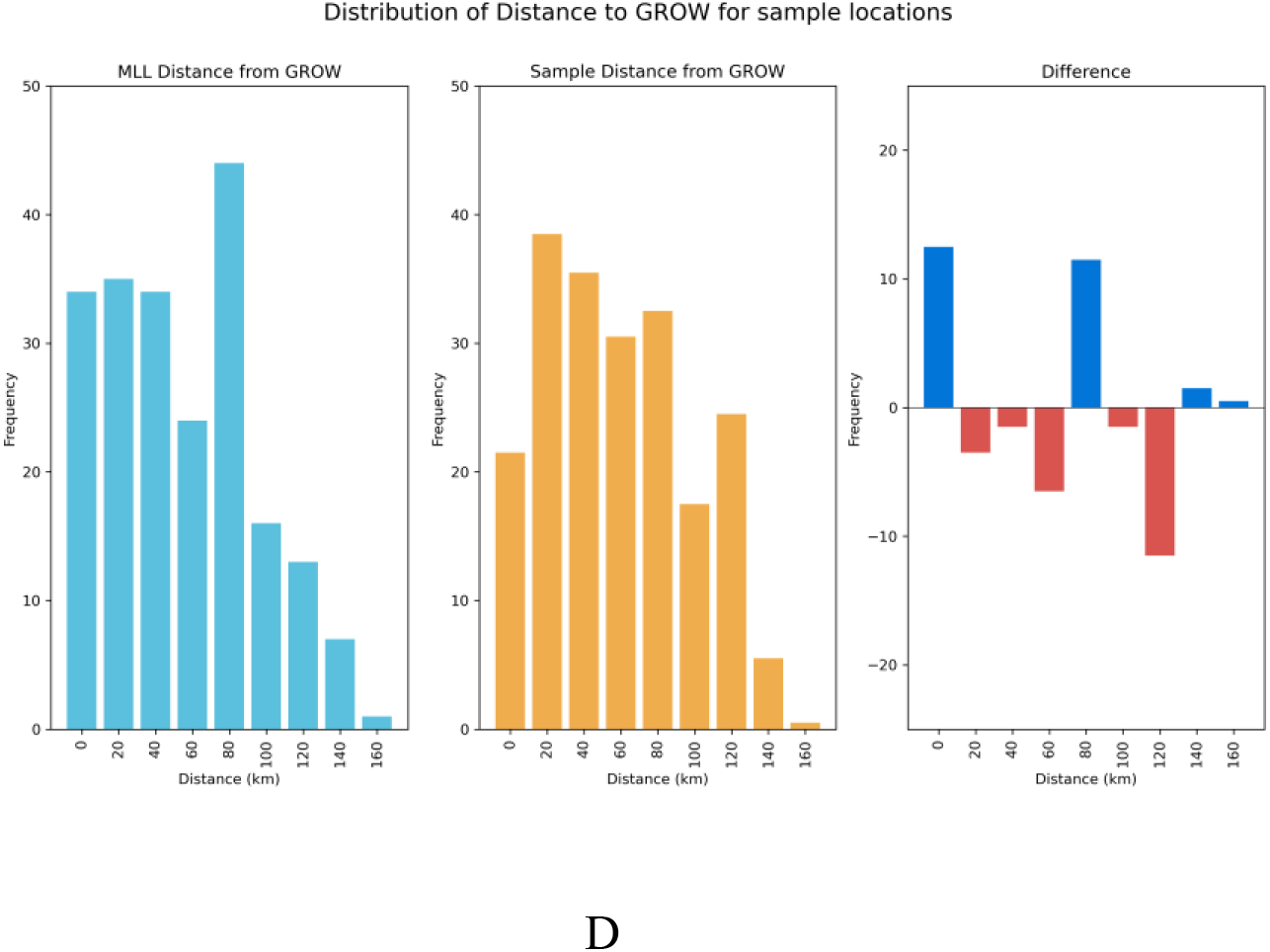
The distribution of MLL victims according to gender (A), Level of education (B), Age (C) and distance they have to travel to gain access to prosthetic services (D)

### The Spatial Distribution of MLL in Acholi

The distribution of those with MLL within the region is shown in Figure 4, aggregated to sub-county level. Spatial autocorrelation analysis using Moran’s *I* statistic [23] does not provide significant evidence of clustering of parishes with a high incidence of MLL (*p*=0.11), suggesting that the MLL is distributed across the entire region, and is not clustered around certain key locations. This is supported by a visual inspection of Figure 4.

**Figure 4:**
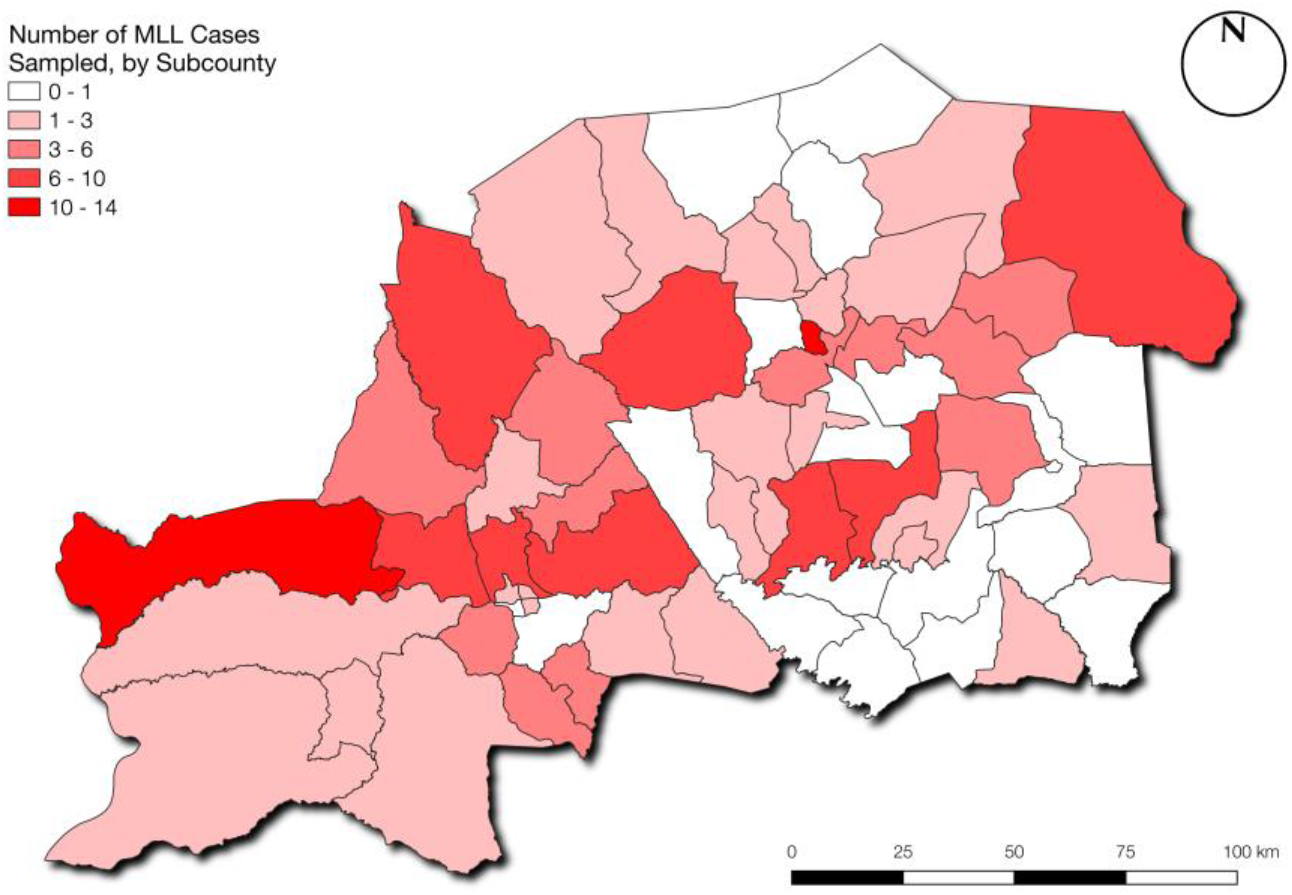
Spatial distribution of MLL victims within the region

Fig. 3d illustrates the observed distribution of distances for the 167 people with MLL for whom parish location was successfully recorded, alongside the expected distribution (created by scaling the distribution of distances of the 800 sample locations, illustrated in Fig. 1). The distances were calculated from the respective parish centre to the Gulu Referral Orthopaedic Clinic, which is the only facility available in the region where prosthetic limb services may be accessed by people with MLL. Whilst the mean and standard distribution of the two distributions are similar (mean: 68.73km; SD: 40.34km expected; mean: 63.67km; SD: 39.66km for those with MLL), the χ^2^ goodness of fit statistic indicates that the probability of the distribution of values having been drawn from the expected distribution is very small (*p*=0.01), and significant at the 99% confidence level. The large positive differences between the observed and expected distributions in the 0-20km and 80-100km bands are explained by the presence of the only settlements of significant size in the region: Gulu, which is the largest town in the region; and Kitgum, about 90km north east of Gulu (Fig. 1).

## Discussion

### Summary of Findings

In this large cross-sectional study, we have demonstrated provided the first systematic estimate of MLL prevalence in the Acholi Sub-Region, despite the fact that the conflict ended in 2005. We estimate that there are approximately 10,117 people with MLL in this region, giving a prevalence rate of approximately 0.5% of the population. We have also demonstrated that those people with MLL in the region are predominantly male, older and have a lower educational attainment when compared to the general population. These results meet with expectations, as males are more likely to have been injured during combat in the LRA insurgency, as well as being more likely to be working in the fields, where they may pick up injuries from remnant land mines or accidents caused during manual labour. The end of the LRA insurgency in around 2005, means that the very young will have experienced a reduced (though not necessarily entirely diminished) effect of the conflict, and are less likely to have been involved in fighting or otherwise become injured due to the conflict (e.g. during an attack or through standing on a landmine, though landmines do still cause injuries amongst children to this day). Those who lived through the conflict will also have likely experienced periods of reduced access to healthcare; meaning that less severe injuries might have resulted in MLL that could otherwise have been treated. It is also perhaps less likely that young people will have developed chronic diseases leading to MLL, such as type-2 diabetes, which typically presents later in life. With respect to levels of education, there is likely a dual causation, with those who are disabled less likely to be able to afford or access education; whereas those with lower levels education are more likely to be in situations where they might become injured, such as engaging in manual labour, working in fields where they might become injured by remnant land mines, or driving a motorbike taxi (boda-boda) and becoming involved in traffic accident.

### Rehabilitative Service Provision

The inadequacy of existing health service provision for those with MLL is illustrated by the spatial analysis presented in this research, which demonstrates that people with MLL are distributed throughout the entire region, as opposed to spatially clustered in certain locations. Based on the current provision of rehabilitation and prosthetic services (a single facility at Gulu Referral Hospital, which is partly funded by this research project), those with MLL would be required to travel an average of over 60km and a maximum of over 170km in order to access these services. The estimated prevalence rate of MLL and this extremely limited service provision, combined with the information presented here about the number of people with MLL who have not had access to assistive devices and rehabilitative services, clearly suggests that the needs of those with MLL are not adequately met by the current level of service provision in this region. War trauma remains the main cause of MLL in our study population and this finding is sharply in contrast with a more recent hospital-based study of 233 amputees where non-communicable diseases accounted for the majority of amputees [24], which reinforces the view that many who suffered MLL in the conflict have not since been able to access medical or rehabilitative support. The inaccessibility of prosthetic and rehabilitative services, high levels of poverty in the region and poor road and transport links suggest that alternative service delivery models are urgently required, as it is extremely unlikely that many subjects will be able to travel to receive services in ‘centralised’ models.

In order to address this issue, we have therefore set up an outreach prosthetic limb service operating from Gulu Referral Hospital as part of this research, in order to demonstrate the potential of such an alternative service delivery model. This outreach service was modelled on a similar approach adopted in post-war Sri Lanka [25], and seeks to deliver prosthetic limb services at open clinics held in local community centres. This is important, as this approach does not require individuals to travel to the clinic and is therefore much more accessible. People with MLL attending the clinic are assessed by a suitably qualified practitioner free of charge, and those with suitable healthy stumps receive bespoke prosthetic limbs free of cost at a location close to their homes. As part of this project, two Ugandan prosthetic technicians also completed a three-month training period at the prosthetic clinics in Sri Lanka, establishing an important South-South collaboration in outreach service delivery models for the rehabilitation of people with MLL in a post-conflict setting. In 2019, this outreach service successfully provided 52 prosthetic limbs to 51 patients (one bilateral). Follow up studies are now needed urgently to further evaluate the potential of such service outreach models, both in terms of efficiency and acceptance by the local population.

### Study Limitations

There are a number of assumptions that had to be made in this research, chief amongst was were the estimate of the population size of the Acholi sub-region, which is a prediction based upon expected population growth from the 2014 census of household population [11], and official refugee numbers [12], both of which are expected to be significant underestimates, and also exclude those living in institutions (e.g. prisons), forest reserves, barracks, and other non-household settings [10]. Further, the 2014 census was based on a sampling (rather than comprehensive) survey, introducing unknown levels of uncertainty in the resulting population figures. This uncertainty is compounded by the difficulty in predicting population change since 2014 due to the extremely rapid population growth and frequent movement of people (including refugees) both into and out of Uganda across adjacent international borders.

The expected distributions for gender and educational levels were based upon data for the Acholi Sub-Region, taken from the 2014 census. We must assume that these distributions (rather than individual values) have not changed significantly in the 4 years between the publication of the census and the data collection, and we are reliant upon the quality of the census data. The expected distribution of ages was based upon the UBOS Mid-Year Population Projection for 2018, which is a modelled extrapolation from the 2014 census. This is a national dataset (values are not given at regional, sub-regional or district level), and so we must assume that the underlying model is robust, and that the population distribution of the Acholi Sub-Region is not significantly different to the national population distribution. The risks associated with all three of these expected distributions are mitigated somewhat by the vanishingly small probabilities of the values in our sample being drawn from the expected distributions, and our deliberate over-sampling is intended to allow for these uncertainties.

The HRSL dataset [16] describes estimated population density, rather than individual buildings. There is also a possibility that this dataset might perform less well in rural than urban settings (as there are fewer buildings available in a more-sparse distribution, which can make less-populated areas harder to detect). However, two ‘on the ground’ validation surveys of the area by the authors, as well as visual validation against high resolution satellite imagery showed that population patterns in the dataset correlate well with the location of buildings and huts in rural areas. Nevertheless, more detailed maps and population data would permit more precise population estimates to be undertaken, and increase the robustness of the sampling strategy.

An error made by some of the trained researchers during data collection meant that precise locations were only collected correctly for 57 of the 181 people with MLL. This was addressed by aggregating MLL sufferers to parish level, for which data were available for 167 people with MLL. Given the relatively small size of the parishes in comparison with the study area, their centroids provide a reasonable proxy location for sample locations, meaning that this has had only a minor impact upon the analysis of distance distribution, and does not affect the quality of the results. Because data were aggregated to sub-county level for the spatial autocorrelation analysis, this limitation has not affected the results of this analysis. Distances from sample locations to the Gulu Referral Orthopaedic Clinic (Fig. 1; Fig, 3d) are given as straight-line distance, which is a significant underestimation of the distance that would have to be travelled by an individual. This is because insufficient map data are available for this region [15], preventing more representative ‘network distances’ (along roads and paths) from being calculated.

### Implications of the Findings

The physical and mental wellbeing of people living in parts of the world that are recovering from prolonged armed conflicts is receiving increasing attention [26, 27]. It is known that people who suffer major disabilities are prone to developing a wide range of secondary health problems as a result of chronic inactivity, social isolation, unemployment (and under-employment), discrimination and marginalisation. For example, a study of landmine victims in Northern Sri Lanka showed that a large proportion of victims lost their earning capacity as a result of their injuries [28]. Furthermore, the prevalence of post-traumatic stress disorder, acute stress reaction, anxiety disorders and depression amongst the victims was significantly greater than in the general population living in the affected areas [28]. These residual issues, attributable to the direct or indirect consequences of the conflict, have a serious impact on quality of life, criminal behaviour, social cohesiveness, productivity and wellbeing. Most importantly, they increase the chances of these communities slipping back into conflict, and thereby trapping many of these already impoverished countries in recurrent cycles of violence and poverty. Though this is a local prevalence study, the methods for undertaking such a study in data poor areas, as well as the outreach clinic model, would be easily transferrable to other settings.

The new insights into MLL and disability provided by this research are vitally important, as a better understanding of the prevalence and nature of MLL will, for the first time, provide an evidence base against which medical care and rehabilitations services can be can be provided to those with MLL in the Acholi sub-region. We have also identified that >8% of the population (>190,500 people) in this region currently live with other forms of major physical disabilities, which may have a major impact on their life chances and quality of life. These findings reinforce the need for greater focus on developing rehabilitation services to meet the needs of the population. War, as anticipated, has been a major cause of MLL in the region, but several other significant causes of MLL have been identified, which cumulatively account for around half of cases, and notably include road traffic accidents and chronic disease. This is critically important in understanding changing service provision needs in this region, as the NGO currently providing these services provide support only to those who lost limbs as a result of the war; meaning that approximately half of the population of people with MLL have no access to rehabilitation services (including prosthetic devices) whatsoever.

The preponderance of lower limb loss over upper limb amputations from combat areas has been reported previously. For example, Bendinelli reported that almost 50% of civilian adults who sustained blast injuries in Cambodia had lower limb injuries [29]. As a result, most prosthetic limb services in post conflict LMICs (including Cambodia, Uganda and Sri Lanka) have focused on lower limb prostheses. Our study revealed that approximately 40% of those with MLL had lost upper limbs, with 2% having lost both upper and lower limbs. Blast injuries from the ground upwards are likely to affect lower limbs more than upper limbs; but amputation of both upper and lower limbs was a widely practiced punishment by the LRA [3], which (along with gunshot wounds) is likely to account for this relatively high proportion of upper limb loss in our study population. The question of rehabilitation for these upper limb amputees is compounded by the absence of any upper limb prosthetic limb services in the Acholi sub-region. Our outreach clinics have also identified many people with MLL that have complex stump-related conditions (protruding bony spikes, infected sinuses, neuromas etc.) that require further corrective surgery before a prosthetic limb can be fitted. Patients with such conditions are offered transport to the Gulu Referral Orthopaedic Clinic, where they can receive the associated surgery prior to being fitted for prosthetic limb.

## Conclusion

This research has provided the first detailed study of disability and major limb loss in the Acholi sub-region, a remote, rural and poor region of Northern Uganda. The information provided in this study, combined with the successful demonstration of an outreach clinic, provides a clear understanding of the level of need for rehabilitative services in the region, as well as a clear agenda for the ways in which these needs can be met through action be researchers, INGOs and government. Further formal studies are now required to evaluate the acceptability, suitability and cost-effectiveness both of outreach rehabilitation services and the prosthetics that we have provided, in order to support their adoption in other post-conflict locations in the world.

## Titles of Figures

**Figure 1.** - Locations of the 8,000 randomly selected sample locations, with an indication of the straight-line distance to the Gulu Referral Orthopaedic Clinic.

**Figure 2.** - Frequencies of causes of MLL in the Acholi sub-region.

**Figure 3.** - Comparison of the observed and expected distribution of (A) gender; (B) education;

(C) age; (D) Euclidean distance to the Gulu Referral Orthopaedic Clinic amongst those with major limb loss.

**Figure 4.** - Locations (by parish) of the 158 MLL sufferers for which valid location data was available, demonstrating a spread of MLL across the whole region

## Supporting Information

**S1.** The Questionnaire administered to 8,000 consenting household heads (S1.household-head-questionnaire.docx).

**S2.** The Questionnaire administered to 181 consenting people with MLL in the surveyed household (S2.mll-questionnaire.docx).

## Conflicts of Interest

The authors certify that there is no conflict of interest with any financial organization regarding the material discussed in the manuscript.

## Funding

This research was funded by the AHRC / MRC GCRF Global Public Health Scheme under Grant AH/R005796/1.

## Author Contributions

Huck led the preparation of the manuscript, which was critically revised by Atim, Moro and Nirmalan. Huck led the statistical and GIS analyses, and wrote the Python code to complete these analyses. Atim prepared an early draft of this manuscript and oversaw the collection and curation of the data. Nirmalan, Moro, Huck and Redmond were responsible for the funding acquisition. Okello and Magada undertook the outreach clinics. All authors contributed to the conceptualisation of this research, and have read and approved the final version of the manuscript.

## Acknowledgements

We thank the Ministry of Health (Uganda) for granting permission to use health facilities for in this study and the district authorities of the 8 Districts and their respective District Health Teams for their support. We would like to thank the *#Huckathon* / Community Mapping (https://communitymapping.org) volunteers in both the UK and Uganda for their continued dedication to mapping the region. The analysis presented here relies upon several Open Source Python libraries, notably Numpy (https://numpy.org), SciPy (https://www.scipy.org) and PySAL (https://pysal.org), GeoPandas (https://geopandas.org), Pandas (https://pandas.pydata.org) and PyProj (https://pypi.org/project/pyproj). Finally, we thank the Acholi People for their kindness, generosity and friendship.

## Footnotes

1 1 arc-second is ^1^/3600 of a degree of latitude or longitude.

## Notes

### Competing Interest Statement

The authors have declared no competing interest.

